# Eco-evolutionary dynamics of pathogen epidemic timing in a seasonal environment

**DOI:** 10.64898/2025.12.08.689079

**Authors:** Ryuichi Kumata, Akira Sasaki

## Abstract

Seasonality structures the dynamics of infectious diseases, yet how pathogens evolve their seasonal timing remains unclear. Here, we develop an evolutionary epidemiology model to examine how pathogens adapt to seasonally varying environments, assuming that each strain favours a specific season for transmission. Analytical and numerical results reveal two contrasting regimes: phenological drift, where the preferred season continually advances through priority effects, and stable seasonal adaptation, where evolution converges to a fixed season. Under strong transmissibility, multiple seasonal morphs can coexist, forming polymorphic communities of epidemic timing. These outcomes result from the balance between a seasonal priority effect, rewarding early occupation of the niche, and a seasonal stabilising effect, adjusting traits to the external optimum. Our findings show that epidemic seasonality itself can evolve as a life-history trait, providing a conceptual bridge between pathogen dynamics and the broader evolution of phenology in periodic environments.

## Introduction

Seasonality is a hallmark of many infectious diseases. Understanding seasonal patterns is crucial for designing interventions, from vaccination calendars to agricultural management (Altizer et al., 2006; Martinez, 2018). Classic explanations for seasonality in epidemiology emphasize external drivers: climatic conditions such as temperature, humidity, and rainfall alter pathogen stability or vector abundance; host behaviour and demography, including school terms in humans or reproductive timing in wildlife, reshape contact patterns; and host physiology, such as annual rhythms in immune and hormonal systems, modulates susceptibility (Arthur et al., 2017; Fisman, 2007; Grassly and Fraser, 2006). These factors together account for familiar patterns such as winter influenza in temperate regions, school-synchronised measles resurgences, rainy-season peaks of malaria and cholera, and agriculturally driven crop diseases (Ferrari et al., 2008; Moriyama et al., 2020; Neumann and Kawaoka, 2022; Reiner Jr et al., 2015; Richard et al., 2022; Shaman et al., 2010). Thus, external seasonality broadly structures epidemic dynamics across taxa.

Yet, external drivers alone do not fully account for the observed diversity of epidemic timing. Pathogens can also evolve intrinsic seasonal preferences that partition ecological niches across the annual cycle. In humans, parainfluenza viruses (HPIV) provide a clear example: HPIV-1 consistently peaks in the fall, whereas HPIV-3 peaks in spring, despite their similar genomes and hosts (Chen et al., 2025; Fry et al., 2006). Other examples of temporal niche partitioning are observed among plant pathogens such as oak powdery mildew and phytophthora fungi (Delmas et al., 2024; Feau et al., 2012; Fodor, 2011; Hamelin et al., 2016). Pathogens may adapt to the season by using cues such as temperature, the host’s immune status, and physiological condition (Martinez-Bakker and Helm, 2015). Taken together, epidemic timing may evolve as a phenological trait of pathogens, determining which part of the annual cycle a pathogen exploits.

Despite the central role of external seasonality in epidemiology, few studies have addressed how pathogens adapt to seasonally structured environments themselves. Previous theoretical work explored how pathogens evolve responsiveness to periodic forcing, showing that selection can favour stronger sensitivity (Kamo and Sasaki, 2005; Koelle et al., 2005). However, these studies focused on the sensitivity of the preference to a specific season rather than the seasonal epidemic timing of pathogens. To our knowledge, no study has addressed the evolution of seasonal preference, the specific seasonal phase at which the pathogen targets to spread.

The evolution of seasonal preference can be viewed within the theory of niche-trait evolution, which explains how organisms differentiate and coexist along resource or environmental gradients through frequency-dependent selection (Dieckmann and Doebeli, 1999; Edwards et al., 2018; Ito and Sasaki, 2023; Leimar et al., 2013; MacArthur, 1970). Previous eco-evolutionary studies have extended this framework to seasonal or periodic environments (Kremer and Klausmeier, 2017; Sakavara et al., 2018). However, these previous studies have included seasonal fluctuations as a factor influencing the adaptation to niche, but did not directly address the evolution of phenological timing. In contrast, this study considers seasonality as the niche to which organisms adapt. A general theoretical framework for phenological evolution in periodic environments remains unexploited (Park and Post, 2022).

Here, we develop a simple evolutionary epidemiology model linking seasonality and adaptation. By representing transmission as a seasonal preference kernel, we show that increasing the amplitude of seasonal fluctuation produces two regimes: phenological drift, where the preferred season continually advances due to priority effects, and stable seasonal adaptation, where pathogens converge to a fixed seasonal niche. We demonstrate that the eco-evolutionary dynamics can lead to polymorphic seasonal patterns in pathogens. These findings illustrate how epidemic seasonality can itself evolve, connecting pathogen phenology to general principles of niche evolution in periodic environments.

## Methods

### Epidemiological model

We model pathogen dynamics using a standard SIRS framework with seasonal forcing in transmission (e.g. Kamo and Sasaki, 2005). Let *S, I*, and *R* denote the fractions of susceptible, infected, and recovered hosts, respectively. We assume that birth and death occur at percapita rate *d* so that the total population size is constant (*S* + *I* + *R* = 1). Recovery from infection and immunity loss occur at rates *γ* and *c*. The above assumption yields the following epidemiological dynamics

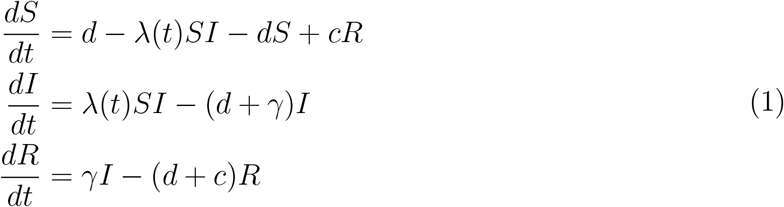

where *λ*(*t*) is the time-dependent effective transmission rate described below.

#### Seasonal phase

We model time *t* measured in year and the seasonal phase *τ* that ranges from zero to one (0 ≤ *τ* ≤ 1) defined as

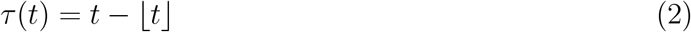

where ⌊*t*⌋ is the floor, integer part, of *t*. For example, at *t* = 7.7, the floor of *t* is 7 and the season 0.7. Notably, phase *τ* is the circular value and *τ* = 0 and *τ* = 1 represent the same phase. All seasonal functions are written in terms of *τ*.

#### Effective transmission rate

We assume that transmission is shaped by two factors: (i) external environmental forcing and (ii) pathogen seasonal preference. Using the seasonal phase *τ*, we specify the time-dependent transmission rate as

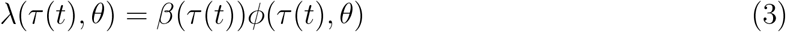

Here, *β*(*τ* (*t*)) represents the seasonal variation in the overall strength of transmission, capturing broad environmental forcing that may arise from host behaviour, environmental variation, or both. In contrast, *ϕ*(*τ* (*t*), *θ*) represents pathogen seasonal preference, specifying the timing within the seasonal cycle at which a pathogen is relatively better adapted. *θ* is the pathogen preferred seasonal phase. Notably, the trait *θ*, representing preferred season, is defined on a circular domain, so that *θ* = 0 and *θ* = 1 correspond to the same seasonal phase. Hence, the relative ordering of phases is not determined by numerical values alone: for example, although *θ* = 0.8 is numerically larger than *θ* = 0, it is more naturally interpreted as lying just before *θ* = 0 on the cyclic niche space.

#### External seasonality

We assume that the environment induces seasonal forcing of transmissibility (Grassly and Fraser, 2006; Kamo and Sasaki, 2005):

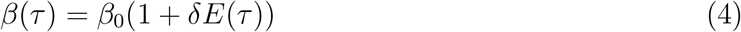

where *E*(*τ*) is the external periodic forcing with period 1 and *δ* is the degree of seasonal fluctuation. We assume the average of external periodic forcing is zero, and the average transmission rate is *β*_0_. In this study, we use a specific function of seasonal forcing *E*(*τ*) = cos(2*π*(*τ* − 1*/*2)) so that transmissibility is maximal at *τ* = 0.5 and minimal at *τ* = 0 (Fig. 1A).

**Figure 1.**
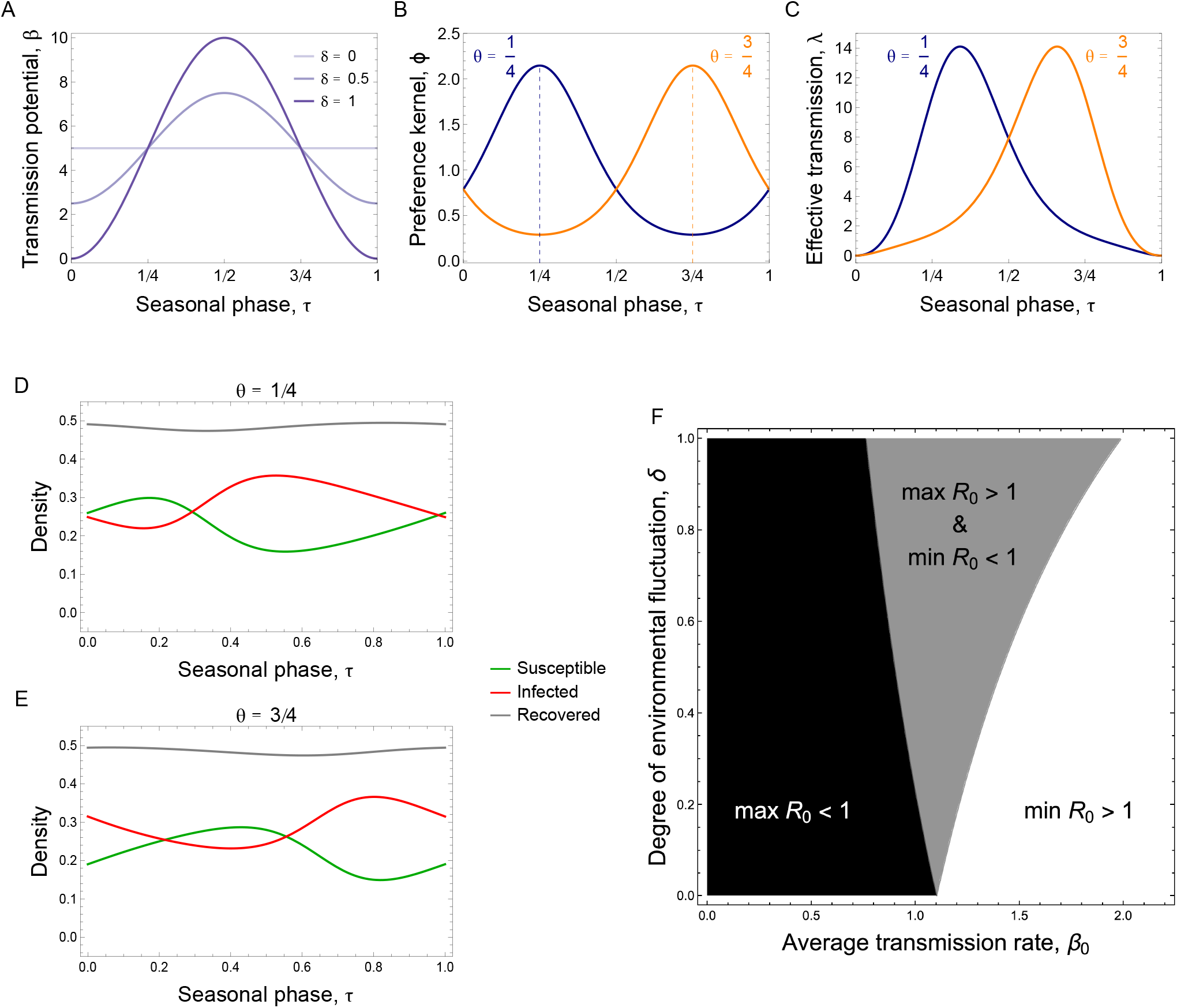
Seasonal preference and seasonal epidemiological dynamics. A. Transmission potential, *β*(*τ*) = *β*_0_(1 + *δE*(*τ*)), which is maximised at *τ* = 1/2. B. Preference kernel *ϕ* with *κ* = 1 and *θ* = 1*/*3(blue) or 2*/*3 (orange). The kernel is distributed over the seasonal phase. C. Effective transmission rate *λ*(*τ, θ*) with *κ* = 1 and *δ* = 1 at the indicated *θ*. D,E. Epidemiological dynamics of pathogens with *θ* = 1/4 (D) and *θ* = 3/4 (E). The trajectory shown here is at the epidemiological attractor. F. Basic reproductive number of the pathogens with seasonal preference. In black area, no strain can cause epidemics. In grey area, some strains can establish epidemics. In white area, all pathogens invade the susceptible population. Parameter values: *d* = 0.1, *γ* = 1, *c* = 0.5, *κ* = 1

#### Seasonal preference kernel

We assume that pathogens express a seasonal preference centred at phase *θ*. We model this preference using a von Mises function

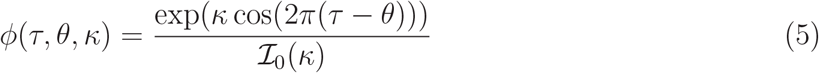

where 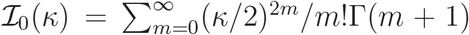 is the modified Bessel function of the first kind, where Γ is the Gamma function. Here *κ* ≥ 0 controls the breadth of the preference, with larger *κ* producing a sharper peak around *θ* (Fig. 1B). This formulation represents a trade-off in seasonal adaptation: greater adaptation to conditions near the preferred phase *θ* comes at the expense of adaptation to other parts of the seasonal cycle. We normalise *ϕ* so that its mean over a year is one, 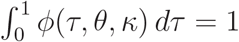. This means that seasonal preference redistributes transmission advantage across the year without changing its annual average. As a result, *κ* controls only the sharpness of seasonal specialisation. Without this normalisation, changing *κ* would also change the annual mean transmission rate, confounding seasonal specialisation with overall transmissibility.

#### Epidemiological dynamics

In this setting, the effective transmission rate, *λ*(*τ, θ*), fluctuates seasonally and peaks when the seasonal environment matches the strain’s preferred phase *θ* (Fig. 1C). This seasonal variation in transmission drives corresponding fluctuations in epidemiological dynamics, and the timing of these fluctuations shifts depending on the value of *θ* carried by the strain (Fig. 1D,E). As the pathogens spread, transmission depletes susceptible hosts, generating epidemiological dynamics that naturally reflect the underlying seasonal pattern of effective transmission.

#### Condition of epidemic

At the disease-free equilibrium (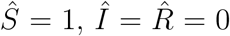), a strain with seasonal preference *θ* can invade if

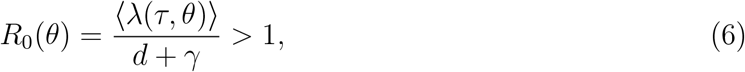

where ⟨·⟩ denotes taking average over a year. In a seasonally varying environment (Eq. (4) with nonzero *δ*), *R*_0_(1/2) and *R*_0_(0) give the maximum and minimum basic reproductive numbers, respectively. We classify the condition of epidemic (Fig. 1C): (i) when *R*_0_(1/2) *<* 1, no strain can invade (black region); (ii) when *R*_0_(1/2) > 1 > *R*_0_(0), only strains near *θ* = 1/2 can cause epidemics (grey region); and (iii) when *R*_0_(0) > 1, all strains having any seasonal preference *θ* can invade (white region). The intermediate regime (grey) emerges only with non-zero seasonal fluctuation in transmission (*δ* > 0).

In the absence of competition between strains, what is optimum for a pathogen is to adjust its preferred season to *θ* = 1/2 where *R*_0_(*θ*) is maximized. However, as shown below, both evolutionary simulations and analytical predictions will show that this optimum will never be attained and diverse evolutionary trajectories will be observed instead.

### Evolutionary simulation

We numerically simulated the evolution of the seasonal preference *θ*. We performed evolutionary simulations using a reaction–diffusion framework on a continuous trait space (Dieckmann and Law, 1996; Kumata and Sasaki, 2022; Sasaki et al., 2022). Importantly, the trait space we considered is cyclic and both ends (*τ* = 0, 1) are connected. In this simulation, the phenotypic distribution of pathogens evolves over time through the epidemiological dynamics (1) and mutation represented by diffusion in trait space *θ*. This approach allows us to track how the distribution of pathogen traits gradually changes and stabilises through the combined effects of selection and mutation.

## Results

### Summary of evolutionary dynamics

We observed that the evolutionary outcome is classified into five distinct patterns (Fig. 2). Figure 2A-E shows the dynamics of trait distribution over time (upper panels) and the density of yearly aggregated *I*(*t*) at indicated years, i.e. slices of the upper panels (lower panels). These patterns are classified based on two criteria: whether the trait distribution is stationary or moving around, and the number of morphs (table in Fig. 2). The first pattern is **mono-drift**, in which a single morph, a localized single-peaked trait distribution, shifts progressively to an earlier phase from year to year (Fig. 2A). The second pattern, **mono-stationary** occurs when the trait distribution converges to a stationary single morph centred at the same phase of the year (Fig. 2B). The third pattern, **multi-drift**, is characterised by a multi-peaked trait distribution that collectively drifts toward earlier seasons (Fig. 2C). In this pattern, multiple morphs coexist dynamically at the evolutionary attractor, forming multiple travelling waves on trait space *θ*. The fourth pattern, **multi-stationary**, consists of a stationary, multi-peaked distribution, indicating the stable coexistence of multiple morphs occupying distinct and fixed seasonal niches (Fig. 2D). The final pattern, **variable**, is defined by the changes in the number of morphs through repeated aggregation and fragmentation of peaks along eco-evolutionary attractor (Fig. 2E). For example, three morphs are present at *t* = 15000, whereas only two remain at *t* = 16500 (lower panel of Fig. 2E). Note that each morph forming a cluster in trait space corresponds to a strain.

**Figure 2.**
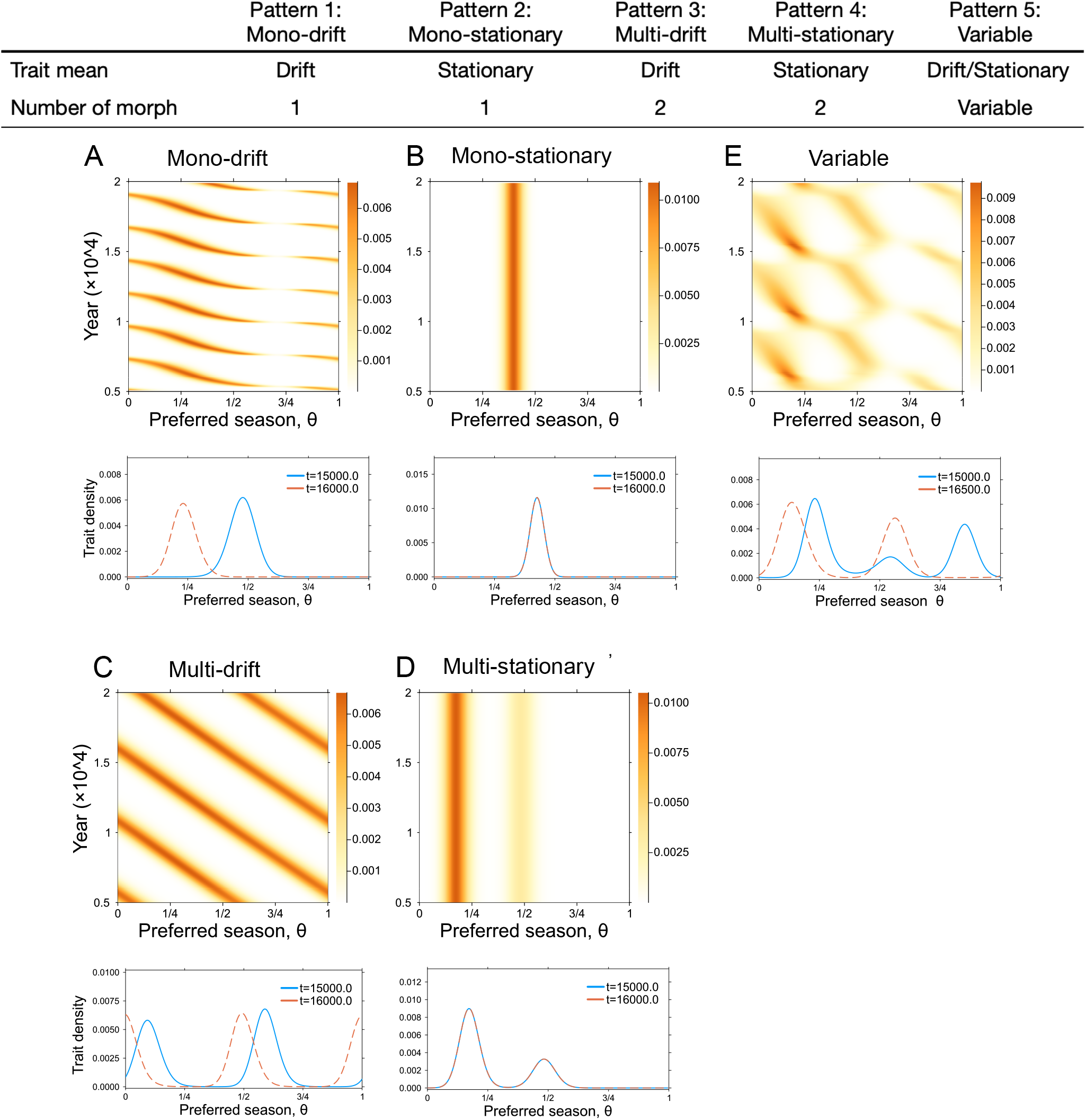
Summary of the evolutionary patterns. The table describes the evolutionary patterns based on two characteristics of trait distribution; trait mean and number of morph. A-E. The distribution of yearly aggregated *I*(*θ*) over 15000 years after burning 5000 years (upper) and the distribution at two indicated time points (bottom) from the numerical evolutionary simulation. Panels A–D are arranged so that columns distinguish drifting versus stationary dynamics, whereas rows distinguish monomorphic versus polymorphic outcomes. Panel E is added separately as an example of the variable regime. We start from a population where most individuals are susceptible, while a small fraction of infected individuals carrying pathogens are randomly distributed, ensuring that the total population sums to one. The trait increment is Δ*θ* = 1*/*200 and the rate of mutation to near traits is *µ* = 0.1 so that the mutational variance is *µ*(Δ*θ*)^2^*/*2 = 1.25 × 10^*−*6^. A *β*_0_ = 2, *δ* = 0.025. B *β*_0_ = 2, *δ* = 0.1. C *β*_0_ = 7, *δ* = 0.01. D *β*_0_ = 5, *δ* = 0.15. E *β*_0_ = 7, *δ* = 0.1. Other parameters are *d* = 0.1, *γ* = 1, *c* = 0.5, *κ* = 1.

### Adaptive dynamical analysis

We first analyse the monomorphic evolutionary dynamics (**mono-drift** and **mono-stationary**). These outcomes are determined by the degree of seasonal fluctuation, *δ*. When *δ* is small, the trait distribution of seasonal preference *θ* forms a sharp peak that gradually shifts forward over time (Pattern 1, Fig. 2A). The population continuously evolves toward earlier seasonal phases, although the evolutionary speed varies depending on the local transmission potential *β*(*τ*). In contrast, when *δ* is large, the trait distribution remains localized around a fixed seasonal niche without further drift (Fig. 2B).

To understand these distinct evolutionary patterns, we analyse the evolution of *θ* based on evolutionary invasion analysis. We consider the fate of a rare mutant invading a resident population on an ecological attractor (Avila and Mullon, 2023; Geritz et al., 1998). Suppose a mutant strain with seasonal preference *θ*_*m*_ appears in the epidemiological equilibrium of a resident strain with preference *θ*. The invasion dynamics of the mutant is

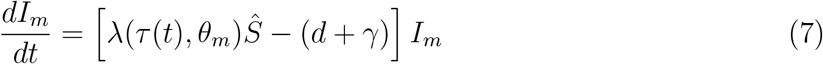

where *Ŝ* = *Ŝ*(*τ* (*t*)) is the density of susceptible hosts at the ecological attractor. In a periodically fluctuating environment, the long-term success of mutant invasions is determined by the timeaveraged invasion fitness over one period (Best and Ashby, 2023; Kamo and Sasaki, 2005; Kremer and Klausmeier, 2017; Lion and Gandon, 2022; Metz et al., 1992). Then, the invasion fitness is defined as

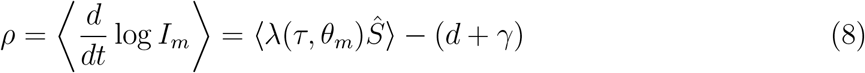

where brackets denote the average over one cycle. The sign of *ρ* determines mutant fate: *ρ >* 0 implies successful invasion, whereas *ρ <* 0 implies failure.

The invasion fitness *ρ* exhibited distinct patterns between the mono-drift and mono-stationary regimes (Fig. 3A,B). In the drift regime, mutants with earlier seasonal preferences had positive fitness relative to the resident, driving a continuous advance of the preferred phase (Fig. 3A). In contrast, in the stationary regime, mutants around the resident trait had negative fitness, indicating evolutionary stability of the resident (Fig. 3B). Pairwise invasibility plots (PIPs; Fig. 3C,D) further illustrate these dynamics (Geritz et al., 1998): when *δ* is small, all residents are invaded by earlier-phase mutants, producing phenological drift; when *δ* is large, two evolutionary singular points emerge—an unstable repeller (grey) and a continuously stable strategy (black) toward which *θ* converges, consistent with simulation results (Fig. 2B).

**Figure 3.**
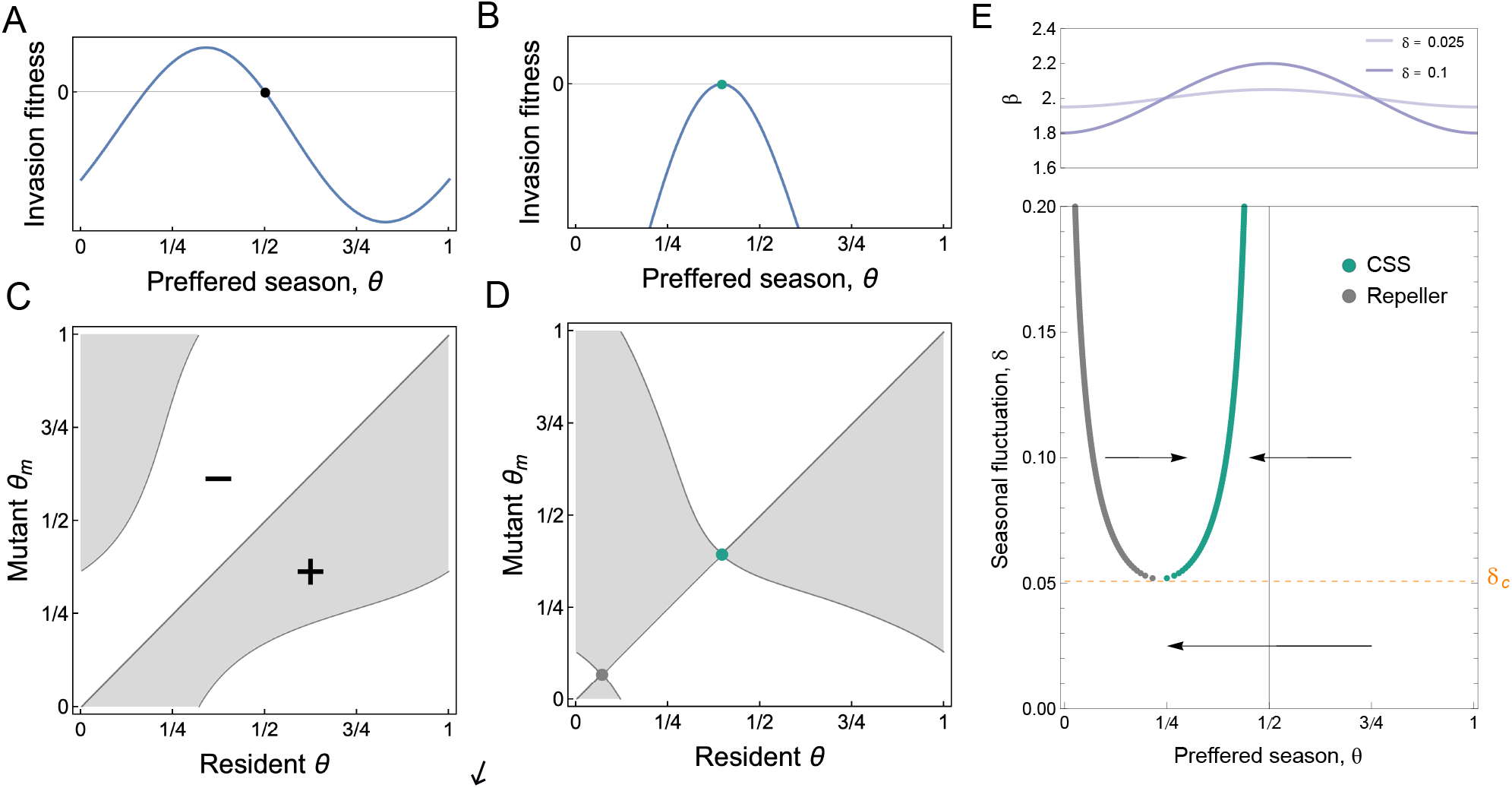
Adaptive dynamical analysis of monomorphic evolutionary dynamics. A, B. Invasion fitness of the mutants when the resident have the indicated phenotype (black and green points). C, D. Pairwise invasibility plot. E. Evolutionary singular points and its convergence stability. Panels A and C represent the case of small fluctuations (*δ* = 0.025), whereas panels B and D represent the case of larger fluctuations (*δ* = 0.1). The upper panel shows the fluctuation of *β* at *δ* = 0.025 and *δ* = 0.1. Parameters: *β*_0_ = 2, *d* = 0.1, *γ* = 1, *c* = 0.5, *κ* = 1

There exists the critical value of degree of seasonal fluctuation, *δ*_*c*_, that separates these monomorphic evolutionary outcomes (Fig. 3E). We mapped the evolutionary singular points for different *δ*’s. When *δ < δ*_*c*_, there is no evolutionary singular points and evolution is always towards earlier seasonal preference (phenological drift). When *δ > δ*_*c*_, an evolutionary repeller and a CSS appear and seasonal preference evolves towards the CSS. A saddle-node bifurcation for evolutionary outcomes occurs at *δ* = *δ*_*c*_. It is important to note that the position of the CSS is always earlier than the environmentally optimal season (*θ* = 1/2) (Fig. 3E).

#### Selection gradient

To interpret the selective force shaping the above evolutionary patterns of the seasonal preference, we focus on the selection gradient 𝒮 (*θ*) of the pathogen’s trait–the preferred season *θ*. This quantity indicates which direction and rate at which *θ* evolves, due to the repeated cycles of invasion and replacement by mutants whose traits slightly differ from the resident (Avila and Mullon, 2023; Geritz et al., 1998).

The selection gradient 𝒮 (*θ*) of the preferred season is defined as the time average of the sensitivity of infection rate *λ*(*τ, θ*) to *θ*, weighted by the susceptible host density *Ŝ*(*τ*):

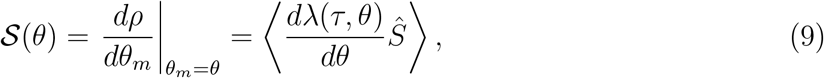

where ⟨·⟩ denotes taking the time average over one seasonal cycle.

By substituting (3) into (9), we can decompose the selection gradient into two components 𝒮_0_ and 𝒮_1_, which corresponds to the contributions from the constant and seasonally varying parts of the environment *β*(*τ*), respectively, where *β*(*τ*) = *β*_0_ + *β*_0_*δE*(*τ*) (details in Supplementary Informaiton):

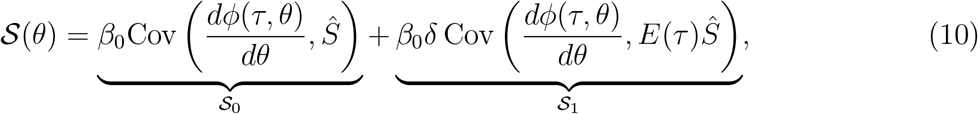

where Cov(*X, Y*) = ⟨*XY*⟩ − ⟨*X*⟩ ⟨*Y*⟩ is the temporal covariance between *X* and *Y* over one period. Note that the time average of *dϕ/dθ* over one cycle is zero (*dϕ/dθ* = 0) owing to the assumed symmetry (Eq. 5).

We first interpret 𝒮_0_. For times *τ* preceding the resident’s preferred season (*τ* < *θ*), *dϕ/dθ* < 0 (see Supplementary Information **S1**), indicating that shifting *θ* earlier would increase infection efficiency during that period. Conversely, for times *τ* following the resident’s preferred season (*τ > θ*), *dϕ/dθ >* 0, meaning that shifting *θ* later increases efficiency during that period. Which of these two opposite effects dominates depends on the sign of the temporal covariance between *dϕ/dθ* and *Ŝ*(*τ*).

Since *Ŝ*(*τ*) is higher before the resident’s preferred season–because more hosts remain uninfected before the infection peak in each year–the covariance Cov(*dϕ/dθ*)(*τ*), *Ŝ*(*τ*)) is negative. This implies that, in the constant environmental part, mutants with an earlier preferred season than the resident are favoured (𝒮_0_ < 0). We refer to this advantage of exploiting uninfected hosts earlier as the “**seasonal priority effect**”.

In contrast, for the component 𝒮_1_ associated with seasonal environmental fluctuation, the direction and strength of selection are determined by the covariance between *dϕ/dθ* and *E*(*τ*)*Ŝ*(*τ*). Because *E*(*τ*)*Ŝ*(*τ*) is positive near the optimum season *τ* = 1/2 and negative near the poorest seasons *τ* = 0 or 1, this component counteracts the seasonal priority effect, pulling the preferred season *θ* back toward the environmental optimum. We refer to this force pulling the preferred season to the environmental optimum as “**seasonal stabilising effect**”.

These two components, the seasonal priority effect (𝒮_0_) and the seasonal stabilising effect (𝒮_1_), have clear ecological interpretations. As shown in Fig. 4A–B, the resident strain progressively depletes susceptible hosts as the season advances, reducing transmission opportunities later in the cycle. A mutant that shifts slightly earlier can therefore infect a larger pool of susceptibles before they are consumed by the resident (Fig. 4A). After replacing the resident, the early mutant becomes the new resident and is again outcompeted by an even earlier mutant, generating a feedback that drives seasonal timing forward (Fig. 4B). This ecological mechanism mirrors priority effects widely observed in community ecology in that reaching a niche earlier gives an advantage (Fukami, 2015; Stroud et al., 2024; Wainwright et al., 2012).

**Figure 4.**
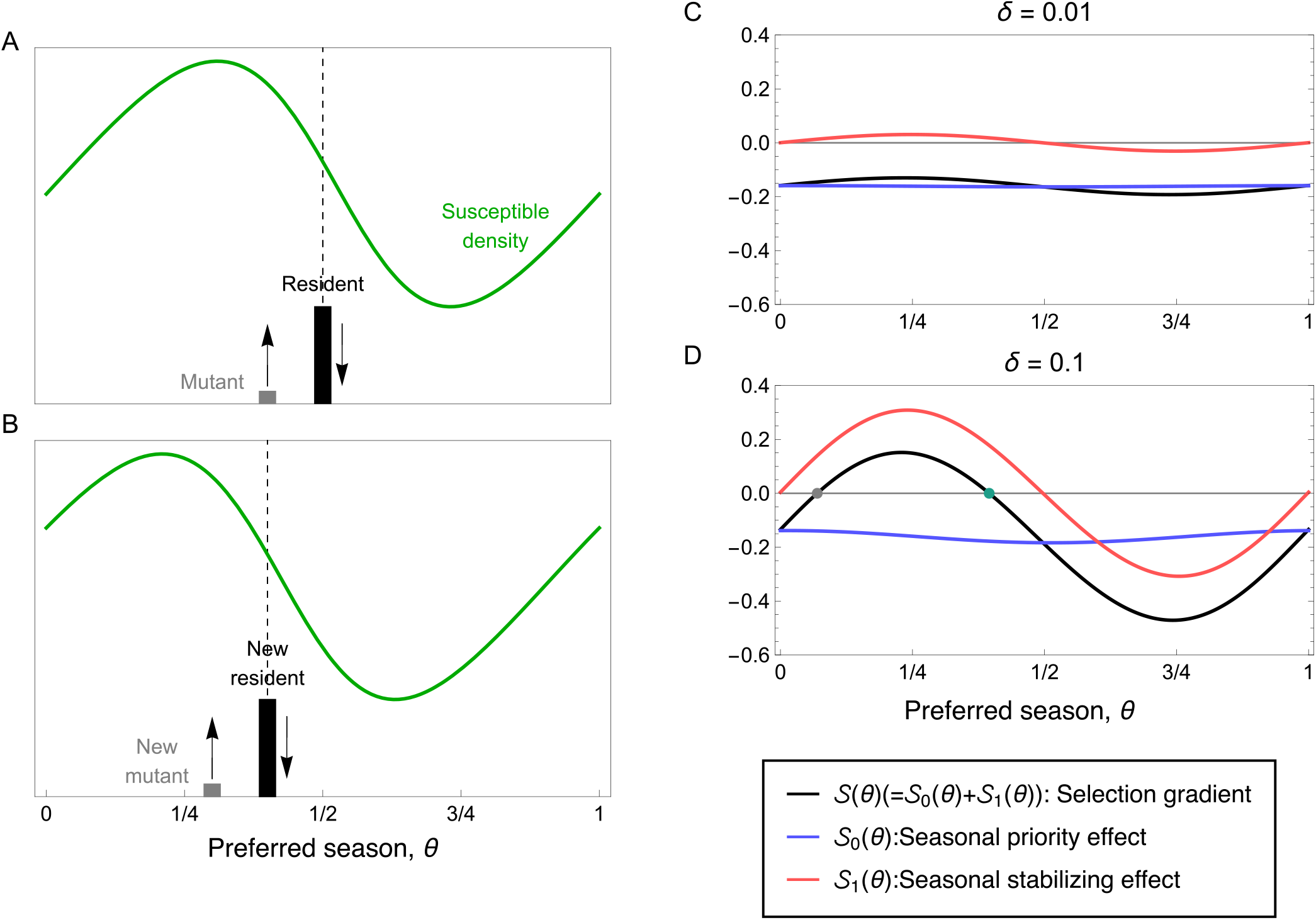
Seasonal priority effect drives the seasonal adaptation. A,B. Illustration for the seasonal priority effect pulling the seasonality to the earlier. Green line is the susceptible density at the attractor *Ŝ*(*τ*) (middle) and black and grey bar represents the resident and the mutant, respectively. C. the selection gradient *S* (black) and its components 𝒮_0_ (blue) and 𝒮_1_ (red). If *δ < δ*_*c*_, 𝒮_1_ is weak and 𝒮_0_ dominates the seasonal adaptation, resulting in the seasonal drift. D. If *δ > δ*_*c*_, 𝒮_1_ is strong enough to make a balance against 𝒮_0_. The black point is the CSS point. Parameters for C, D: *β*_0_ = 2, *d* = 0.1, *γ* = 1, *c* = 0.5, *κ* = 1

By contrast, the second term, 𝒮_1_, captures the cost of deviating from the timing at which external environmental conditions most strongly promote transmission. Because this component is proportional to the amplitude of seasonality *δ*, it becomes influential when environmental fluctuations are pronounced. 𝒮_1_ increases the fitness of strains whose preferred season aligns with the seasonal peak of transmission opportunities (*τ* = 0.5 in our assumption) and imposes costs on shifts away from this optimum. Thus, 𝒮_1_ represents the seasonal stabilising effect, which pulls *θ* toward the environmental optimum.

The balance between these two forces determines whether evolution produces continual advancement or settles at a continuously stable strategy (CSS). When seasonality is weak (Fig. 4C, *δ* = 0.01), 𝒮_1_ is negligible and the dynamics are dominated by 𝒮_0_, which remains negative across all *θ*. This imbalance drives a continual evolutionary shift toward earlier seasonal preference. When seasonality becomes stronger (Fig. 4D, *δ* = 0.1), 𝒮_1_ increases in magnitude and introduces a stabilising force that counteracts 𝒮_0_ at a particular value of *θ*. This balance produces a continuously stable strategy (CSS), where directional and stabilising forces are in equilibrium. The detailed description of these effects is summarised in Supplementary Information S1. Taken together, the evolutionary trajectory of seasonal preference is governed by seasonal priority effect and seasonal stabilising effect.

We also found that some evolutionary singular points are convergence stable but evolutionarily unstable (Fig. 5A). Such points correspond to potential branching points, where a monomorphic population is expected to diversify through disruptive selection (Avila and Mullon, 2023; Geritz et al., 1998). We then classified the stability of evolutionary singular points and identified the threshold value *δ*^*^ that separates the stability (Fig. 5B). When the baseline transmission rate *β*_0_ is smaller than a value *δ*^*^ is overlapped by *δ*_*c*_ (orange line). When *β*_0_ is larger than a value (around six), the threshold *δ*^*^ is a slightly greater than *δ*_*c*_. Notably, evolutionary branching predicted by AD occurs only in a narrow parameter region characterised by *β*_0_ and values of *δ* in the range from *δ*_*c*_ to *δ*^*^ (Fig. 5B).

**Figure 5.**
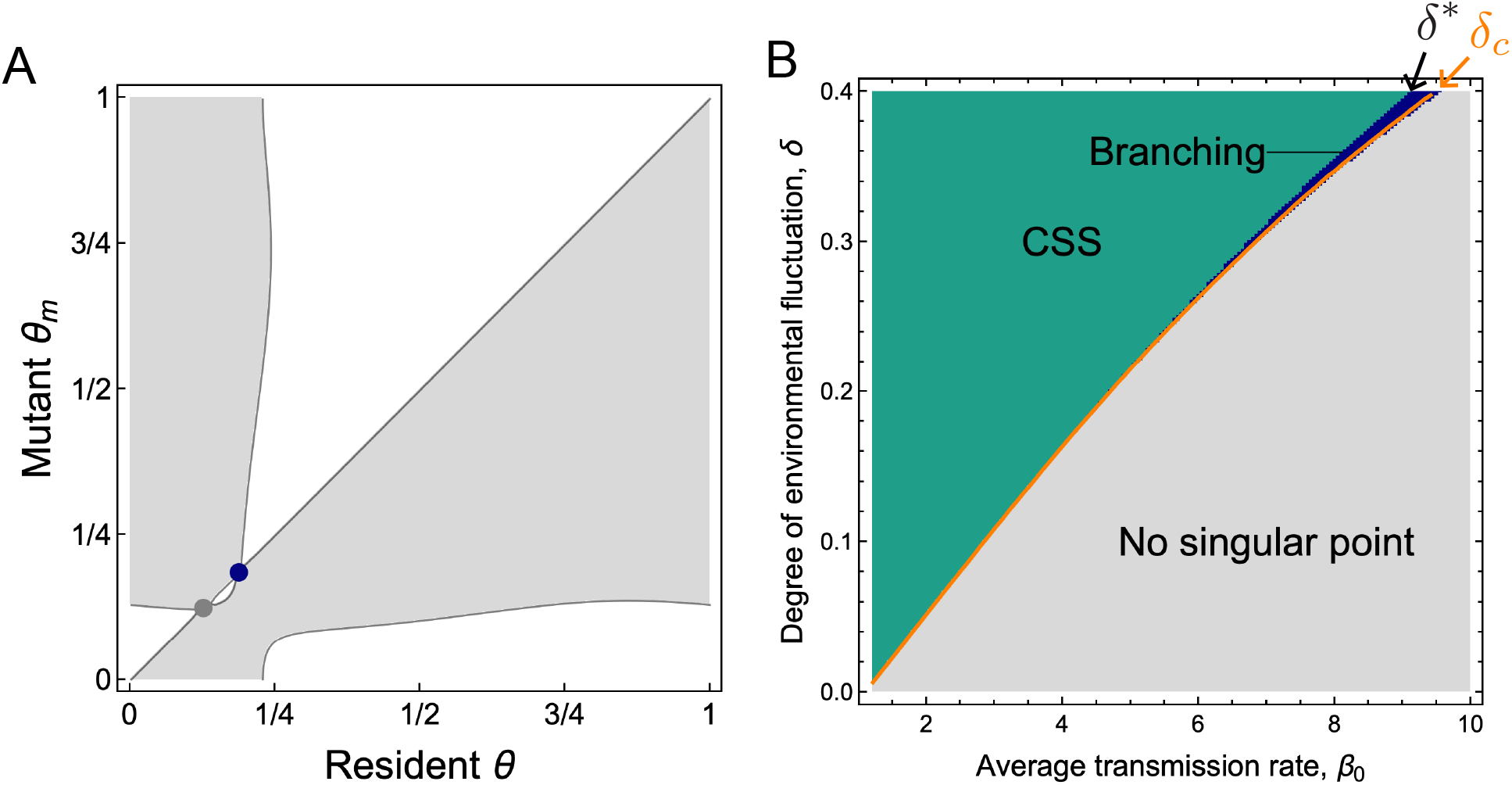
Evolutionary stability of singular points. A. PIP indicates evolutionary branching. Dark blue point is convergence stable but evolutionarily unstable. B. Stability of singular point. Dark green area shows the area where there is a single CSS point and a repeller point. Light grey area does not have any singular point. Dark blue area is the area where the convergence stable point is evolutionarily unstable. Orange line is *δ*_*c*_ obtained from AD analysis (Fig. 3E). Parameters: *d* = 0.1, *γ* = 1, *c* = 0.5, *κ* = 1 (for both), and *β*_0_ = 9.35, *δ* = 0.4 (for A)

However, the multi-morphic evolutionary dynamics are frequently observed in our numerical evolution. Indeed, the complex polymorphism dynamics observed in our simulations (Figs. 2C–E) arise precisely outside the regions of evolutionary branching (e.g., Figs. 2D with *β*_0_ = 5, *δ* = 0.15). Consequently, the polymorphic evolutionary dynamics occur more robustly in our simulations than predicted by the evolutionary branching condition of AD. Furthermore, our simulations revealed that the critical transition between monomorphic drift and monomorphic stationary pattern strongly depends on mutational variance (Fig. S2). This dependence cannot be captured by the classical AD framework, which assumes infinitesimal mutational variance and tracks only the mean trait. These results emphasise the need for analytical frame-works beyond AD that explicitly consider finite mutation variance and multiple phenotypes.

### Oligomorphic dynamical analysis

To overcome this limitation and fully capture the parameter dependence of evolutionary outcomes, we next adopt the oligomorphic dynamics (OMD) framework (Lion et al., 2023; Sasaki and Dieckmann, 2011). OMD is a framework that extends adaptive dynamics by treating each of multiple coexisting morphs as having a narrow, Gaussian-shaped trait distribution (Fig. 6A). Based on this approximation, one can derive the coupled dynamics of ecological densities, morph frequencies, and the key moments of the trait distribution within each morph (its mean and variance). OMD yields the evolutionary trajectory of each morphs, enabling us to capture the evolutionary patterns that is realised under a set of given parameters. We derived a set of equations of OMD in our model (Supplementary Information S3) to investigate the evolutionary patterns beyond the scope of AD approach.

**Figure 6.**
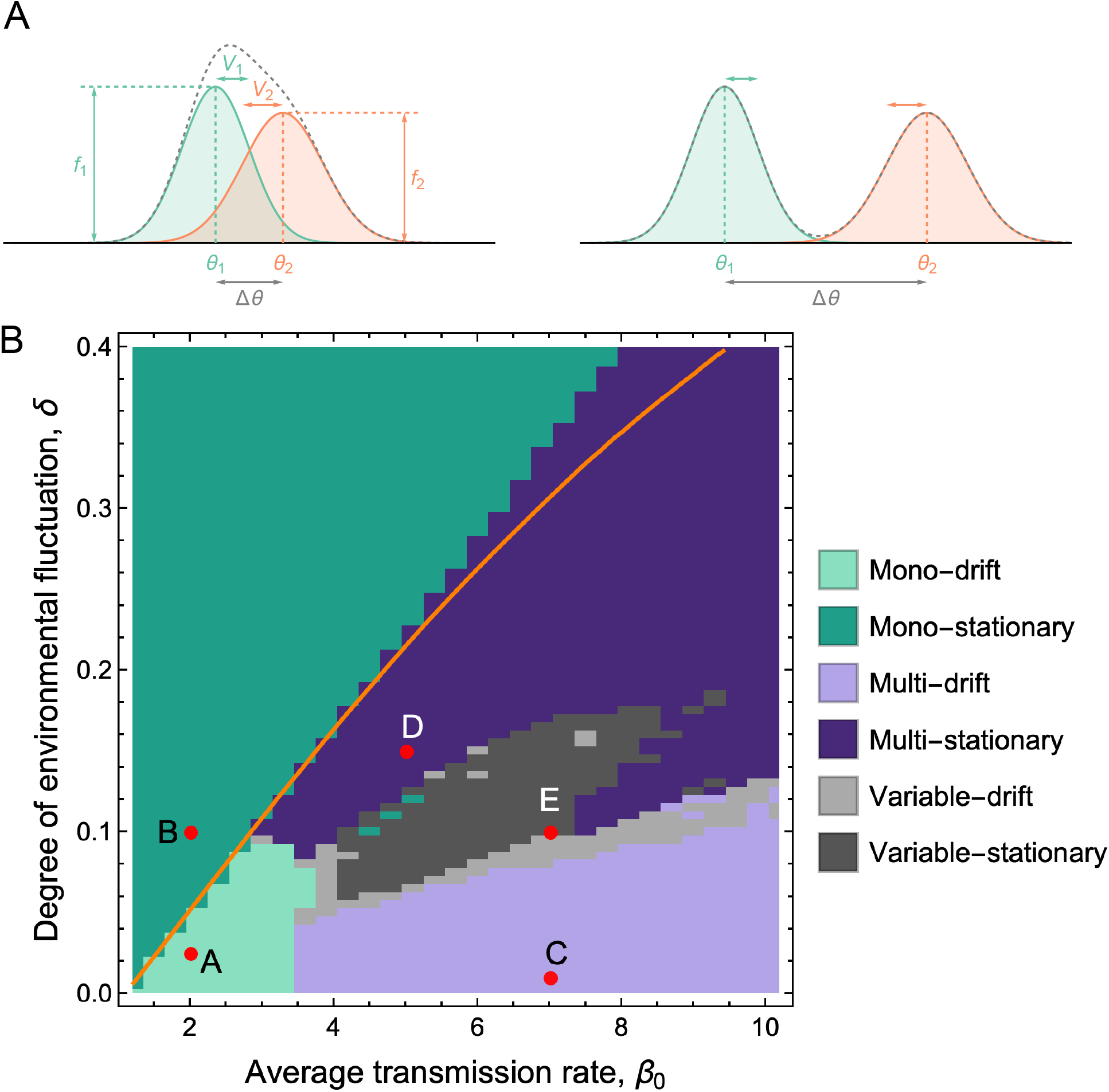
Prediction of the evolutionary patterns by OMD. A. Illustration of oligomorphic assumption. The full distribution (dashed line) is composed of multiple Gaussian distribution with mean *θ*_*i*_ and variance *V*_*i*_ weighted by frequency *f*_*i*_. When two morphs are closer than a threshold (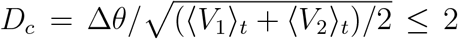) (**Box 1**), two morphic distributions are largely overlapped and the full trait distribution (dashed line) is judged as a single morphic distribution. In contrast, when two morphs are apart (*D*_*c*_ > 2), the full distribution is judged as a composition of multiple morphs. B. The evolutionary pattern predicted by OMD. Numerical method is described in Supplementary Information S4. Lighter colours indicate the drifting behaviour, while darker colours indicate that the distribution is fixed or fluctuating around a set of specific seasons. Orange line is *δ*_*c*_ obtained from AD analysis (the same in Fig. 5B). Each red point is the corresponding points shown in Fig. 2A-E. Parameters:*d* = 0.1, *γ* = 1, *c* = 0.5, *κ* = 1, *V*_*m*_ = 0.1 * 0.5 * (1/200)^2^/(2*π*)^2^

We found that OMD captures the dependency of the critical transition *δ*^*^ on mutational variance (Fig. S2). Larger mutational variance lowers *δ*^*^, making it easier for the system to reach a stationary distribution. One potential explanation for this pattern is that broad standing trait variation generated by large mutational variance spread transmission events over a longer part of the seasonal cycle. This can also stretch out the period during which susceptible hosts are depleted. As a result, the directional selection gradient becomes weaker, making it hard to maintain the drifting when variance is large.

OMD gives the comprehensive prediction of evolutionary patterns including the regime with no evolutionary singular points by AD analysis (Fig. 6B). The numerical scheme for classifying the evolutionary dynamics is summarised in Summplemetary Information S4. Multi-morphic patterns are generated when *β*_0_ is greater than around 3.5. A high transmission rate accelerates the speed of epidemiological dynamics and lowers the threshold for the susceptible density required for an epidemic to become possible. Consequently, it is expected that after the epidemics of a preceding strain, another strain may be able to spread again within a year. This is likely the logic maintaining polymorphism through high infectivity. Furthermore, as with the monomorphic case, the degree of seasonal fluctuation impacts whether the evolutionary pattern is drifting or not. When *δ* is small (light purple area), **multi-drift** is observed, whereas sufficiently large *δ* generates **multi-stationary** pattern. Between these two regimes, the evolutionary pattern is predicted as **variable**, in which the number of morphs changes along time. In polymorphic populations, increasing *δ* imposes heterogeneous selective pressures across morphs because each morph is associated with a different seasonal phase. In this intermediate region, these morph-specific selective forces are strong enough to prevent seamless drifting, but not yet strong enough to stabilise a fixed number of morphs at fixed trait positions, resulting in complex non-stationary dynamics. The upper boundary of the multi-morphic region lies close to *δ*^*^, the bifurcation point predicted by AD (orange line). These classifications are also consistent with the results of evolutionary simulations: each red point corresponds to the numerical simulation shown in Fig. 2A–E. Furthermore, similar qualitative evolutionary patterns were obtained when other parameters (*d* and *c*) were varied. This demonstrates the validity and robustness of our OMD-based prediction.

## Discussion

Our findings reveal that epidemic seasonality can itself evolve through eco-evolutionary feed-backs between pathogen evolution and seasonal epidemiological dynamics. Traditionally, seasonal timing of pathogens has been attributed mainly to external environmental drivers. In contrast, we treat seasonal preference as an evolving pathogen trait and explore its long-term evolutionary dynamics. Our analysis identified two effects: a stabilising effect imposed by external seasonality acting on transmission rate, and a seasonal priority effect, in which pathogens that initiate epidemics earlier gain a competitive advantage. The evolutionary outcome depends on the balance between these two forces. When external seasonal forcing is weak, the priority effect drives directional selection, leading to evolutionary drift in seasonal preference—implying that epidemic schedules may not be fixed but can cycle evolutionarily. When stabilising forces dominate, pathogens evolve toward a specific seasonal preference, yet it is evolutionarily stabilised at a time earlier than the environmentally optimal time. This reflects a balance between the environmental optimum and the priority effect.

The seasonal priority effect is consistent with short-term observations in infectious diseases. For instance, in respiratory infections such as influenza and RSV, the epidemic size of one strain or subtype is often reduced when another has already circulated earlier in the same season (Nickbakhsh et al., 2019). This phenomenon, known as epidemiological interference, is typically attributed to cross-immunity or temporary depletion of susceptible hosts caused by the earlier epidemic (Bhattacharyya et al., 2015; Nickbakhsh et al., 2019; Rohani et al., 2003). Our model assumes complete cross-immunity between strains, which is consistent with such observations and theoretical assumptions. In U.S. non-polio enteroviruses, the relatively recently emerged CV-A6 lineage shows a distinctly earlier seasonal pattern than other serotypes. This pattern might be related to some evolutionary change driven by a seasonal priority effect (Pons-Salort et al., 2018). Our results thus highlight the possibility that these short-term competitive interactions may scale up to shape the long-term evolution of epidemic phenology and seasonal niche partitioning in infectious diseases (Rohani et al., 2003).

Analogous priority effects are also widespread in broader ecological and evolutionary contexts. In cyclically fluctuating environments, genotypes that exploit resources earlier gain a competitive advantage, creating asymmetrical competition along the temporal niche axis (Matsuda and Abrams, 1994). In plant communities, for example, invasive species with earlier germination often suppress the recruitment of later-emerging natives through priority effects during seedling establishment (Blackford et al., 2020; Cleland and Wolkovich, 2024; Wainwright et al., 2012). Importantly, in the context of evolution regarding seasonal timing, such priority effects are unavoidable, because resident types always compete with mutants over the seasonally distributed resources. Overall, the seasonal priority effects may represent a general evolutionary principle governing the phenology across species.

The intensity of environmental fluctuations impacts evolutionary patterns (Fig. 6). This suggests that the seasonality of pathogens may change due to the effects of climate change: if climate change reduces the magnitude of environmental fluctuations affecting pathogens, the seasonal timing of pathogens may shift earlier. Therefore, understanding how climate change impacts the seasonal component of pathogens transmission is crucial not only for short-term outbreak prediction but also for examining long-term evolutionary dynamics (Baker et al., 2022).

We also observed multi-morphic evolutionary dynamics in seasonal environments, similar to the alternating seasonal peaks of closely related HPIV subtypes and plant pathogens (Delmas et al., 2024; Fry et al., 2006; Hamelin et al., 2011). In our model, the coexistence of multiple seasonal-preference morphs may arises through a simple mechanism: high level of basal transmission enables repeated re-invasion of a pathogen strain after a single epidemic. Although Hamelin et al. (2011) considered phenological diversification of plant pathogens through a more complex life-history trade-off between within-season transmission and between-season survival, our results highlight that the seasonal polymorphism can evolve purely through adaptation to the seasonal environment itself without the life-history details. Taken together, these findings suggest that cyclic temporal structure alone can be a powerful driver of multi-morph coexistence in pathogens, providing the new insight for the understanding of the eco-evolutionary mechanism of coexistence in fluctuating environments (Yamamichi et al., 2023).

The coexistence of multiple morphs links our analysis to species packing theory in fluctuating environments. Classical species packing theory examines how many species can coexist along a linear niche axis under constant, random, or periodic fluctuations (Leimar et al., 2013; MacArthur, 1970; Sakarchi and Germain, 2025; Sakavara et al., 2018; Sasaki, 1997; Sasaki and Ellner, 1995). In our model, the evolving trait represents a seasonal preference on a circular niche axis, where trait values wrap around the annual cycle. On such a periodic trait space, evolution can move persistently in one direction while eventually returning to the same trait state. Consequently, we find not only stable clusters of morphs with different preferred seasons, but also dynamic coexistence of multiple morphs, in which each morphs is moving over the cyclic niche space. Our results therefore extend species packing theory to intrinsically periodic niche structures, showing that fluctuating environments can maintain diversity through dynamic rather than static forms of niche differentiation.

We revealed that oligomorphic dynamics (OMD) theory provides comprehensive predictions of long-term evolutionary outcomes (Fig. 6). Compared to adaptive dynamics (AD), OMD offers several advantages. First, OMD enables us to investigate the effects of variance that are ignored in AD. In particular, OMD captures the dependency of evolutionary dynamics on mutational variance (Fig. S2). Second, OMD enables us to predict complex evolutionary behaviours such as drifting of multimorph distribution, which are beyond the scope of AD. Third, it is computationally more efficient: analysing the existence of static dimorphism requires AD to perform exhaustive invasion analyses for all combinations of resident and variant traits, whereas OMD requires solving only a single system of ordinary differential equations. This advantage is particularly decisive in periodic environments, where invasion fitness must be computed through time integration. To our knowledge, this computational benefit of OMD has not been emphasized in earlier studies. Our study emphasises that OMD provides an excellent computational technique for analysing highly complex evolutionary dynamics, such as those in seasonal niches, which conventional AD cannot handle (Lion et al., 2023; Sasaki and Dieckmann, 2011).

The magnitude of seasonal forcing in transmission potential is likely to depend on both host behaviour and the external environment (Grassly and Fraser, 2006). It should be larger when host contact patterns vary strongly over the year, and also in regions where climatic conditions such as temperature and humidity vary strongly across seasons. Although such effects can be inferred indirectly from epidemiological time series using seasonally forced transmission models (Goeyvaerts et al., 2015), these approaches usually recover only an overall seasonal effect rather than a unique mechanism. Available comparative studies further suggest that HPIV is often less sharply seasonal than RSV, and in some settings less seasonally concentrated than influenza. In Spain, RSV and influenza showed clearly defined seasons, whereas HPIV showed less clear seasonality (Shirreff et al., 2024). In addition, parainfluenza epidemics were reported to last longer on average than those of RSV or influenza (Li et al., 2019). This is consistent with our finding that weaker seasonal forcing favours multi-morphic dynamics, and may help explain why HPIV shows diverse seasonal patterns across viral types and settings (Chen et al., 2025).

Our study assumed that pathogens can adapt to the specific season, but the underlying mechanisms remain poorly understood. One possibility is that pathogens exploit the host’s seasonal clock: hosts often show seasonal cycles in immunity, metabolism, and behaviour (Bradshaw and Holzapfel, 2007; Forrest and Miller-Rushing, 2010; Martinez-Bakker and Helm, 2015), and by responding to these cycles, pathogens may indirectly track specific seasonality. Our analysis also assumed that seasonality acts solely on pathogen transmission, whereas in reality seasonal variation may also arise from host demography and behaviour, such as seasonal activity or breeding patterns (Altizer et al., 2006). Extending our framework to include explicit mechanisms of seasonal sensing and periodicity in host life history is an important challenge for future work.

To conclude, we found a new evolutionary logic determining seasonal timing of pathogens: seasonal priority effect, which yields asymmetric resource competition over the seasonal niche. This principle may provide a general evolutionary foundation for the adaptive evolution of phenology across diverse taxa beyond pathogens.

## Acknowledgement

The authors thank S. Gandon, S. Lion, and two anonymous reviewers for valuable comments on the earlier draft.

## References

Altizer, S., A. Dobson, P. Hosseini, P. Hudson, M. Pascual, and P. Rohani (2006). “Seasonality and the dynamics of infectious diseases”. In: Ecology letters 9.4, pp. 467–484.

Arthur, R. F., E. S. Gurley, H. Salje, L. S. Bloomfield, and J. H. Jones (2017). “Contact structure, mobility, environmental impact and behaviour: the importance of social forces to infectious disease dynamics and disease ecology”. In: Philosophical Transactions of the Royal Society B: Biological Sciences 372.1719, p. 20160454.

Avila, P. and C. Mullon (2023). “Evolutionary game theory and the adaptive dynamics ap-proach: adaptation where individuals interact”. In: Philosophical Transactions of the Royal Society B 378.1876, p. 20210502.

Baker, R. E., A. S. Mahmud, I. F. Miller, M. Rajeev, F. Rasambainarivo, B. L. Rice, et al. (2022). “Infectious disease in an era of global change”. In: Nature reviews microbiology 20.4, pp. 193–205.

Best, A and B Ashby (2023). “How do fluctuating ecological dynamics impact the evolution of hosts and parasites?” In: Philosophical Transactions of the Royal Society B 378.1873, p. 20220006.

Bhattacharyya, S., P. H. Gesteland, K. Korgenski, O. N. Bjørnstad, and F. R. Adler (2015). “Cross-immunity between strains explains the dynamical pattern of paramyxoviruses”. In: Proceedings of the National Academy of Sciences 112.43, pp. 13396–13400.

Blackford, C., R. M. Germain, and B. Gilbert (2020). “Species differences in phenology shape coexistence”. In: The American Naturalist 195.6, E168–E180.

Bradshaw, W. E. and C. M. Holzapfel (2007). “Evolution of animal photoperiodism”. In: Annu. Rev. Ecol. Evol. Syst. 38.1, pp. 1–25.

Chen, J., S. Deng, X. Xu, S. Chen, Y.-N. Abo, Q. Bassat, et al. (2025). “Regional and type-specific variations in the global seasonality of human parainfluenza viruses and the influence of climatic factors: a systematic review and meta-analysis”. In: The Lancet Global Health 13.8, e1425–e1435.

Cleland, E. E. and E. Wolkovich (2024). “Effects of phenology on plant community assembly and structure”. In: Annual Review of Ecology, Evolution, and Systematics 55.1, pp. 471–492.

Delmas, C. E., M.-O. Bancal, C. Leyronas, M.-H. Robin, T. Vidal, and M. Launay (2024). “Monitoring the phenology of plant pathogenic fungi: why and how?” In: Biological Reviews 99.3, pp. 1075–1084.

Dieckmann, U. and M. Doebeli (1999). “On the origin of species by sympatric speciation”. In: Nature 400.6742, pp. 354–357.

Dieckmann, U. and R. Law (1996). “The dynamical theory of coevolution: a derivation from stochastic ecological processes”. In: Journal of mathematical biology 34.5, pp. 579–612.

Edwards, K. F., C. T. Kremer, E. T. Miller, M. M. Osmond, E. Litchman, and C. A. Klausmeier (2018). “Evolutionarily stable communities: a framework for understanding the role of trait evolution in the maintenance of diversity”. In: Ecology letters 21.12, pp. 1853–1868.

Feau, N., A. Lauron-Moreau, D. Piou, B. Marçais, C. Dutech, and M.-L. Desprez-Loustau (2012). “Niche partitioning of the genetic lineages of the oak powdery mildew complex”. In: Fungal Ecology 5.2, pp. 154–162.

Ferrari, M. J., R. F. Grais, N. Bharti, A. J. Conlan, O. N. Bjørnstad, L. J. Wolfson, et al. (2008). “The dynamics of measles in sub-Saharan Africa”. In: Nature 451.7179, pp. 679–684.

Fisman, D. N. (2007). “Seasonality of infectious diseases”. In: Annual review of public health 28.1, pp. 127–143.

Fodor, E. (2011). “Ecological niche of plant pathogens”. In: Annals of Forest Research 54.1, pp. 3–21.

Forrest, J. and A. J. Miller-Rushing (2010). Toward a synthetic understanding of the role of phenology in ecology and evolution.

Fry, A. M., A. T. Curns, K. Harbour, L. Hutwagner, R. C. Holman, and L. J. Anderson (2006). “Seasonal trends of human parainfluenza viral infections: United States, 1990–2004”. In: Clinical infectious diseases 43.8, pp. 1016–1022.

Fukami, T. (2015). “Historical contingency in community assembly: integrating niches, species pools, and priority effects”. In: Annual review of ecology, evolution, and systematics 46.1, pp. 1–23.

Geritz, S. A., E Kisdi, G. Meszé NA, and J. A. Metz (1998). “Evolutionarily singular strategies and the adaptive growth and branching of the evolutionary tree”. In: Evolutionary ecology 12.1, pp. 35–57.

Goeyvaerts, N., L. Willem, K. Van Kerckhove, Y. Vandendijck, G. Hanquet, P. Beutels, et al. (2015). “Estimating dynamic transmission model parameters for seasonal influenza by fitting to age and season-specific influenza-like illness incidence”. In: Epidemics 13, pp. 1–9.

Grassly, N. C. and C. Fraser (2006). “Seasonal infectious disease epidemiology”. In: Proceedings of the Royal Society B: Biological Sciences 273.1600, pp. 2541–2550.

Hamelin, F. M., A. Bisson, M.-L. Desprez-Loustau, F. Fabre, and L. Mailleret (2016). “Temporal niche differentiation of parasites sharing the same plant host: oak powdery mildew as a case study”. In: Ecosphere 7.11, e01517.

Hamelin, F. M., M. Castel, S. Poggi, D. Andrivon, and L. Mailleret (2011). “Seasonality and the evolutionary divergence of plant parasites”. In: Ecology 92.12, pp. 2159–2166.

Ito, H. C. and A. Sasaki (2023). “The Adaptation Front Equation Explains Innovation-Driven Taxonomic Turnovers and Living Fossilization”. In: The American Naturalist 202.6, E163– E180.

Kamo, M. and A. Sasaki (2005). “Evolution toward multi-year periodicity in epidemics”. In: Ecology Letters 8.4, pp. 378–385.

Koelle, K., M. Pascual, and M. Yunus (2005). “Pathogen adaptation to seasonal forcing and climate change”. In: Proceedings of the Royal Society B: Biological Sciences 272.1566, pp. 971– 977.

Kremer, C. T. and C. A. Klausmeier (2017). “Species packing in eco-evolutionary models of seasonally fluctuating environments”. In: Ecology Letters 20.9, pp. 1158–1168.

Kumata, R. and A. Sasaki (2022). “Antigenic escape is accelerated by the presence of immuno-compromised hosts”. In: Proceedings of the Royal Society B 289.1986, p. 20221437.

Leimar, O., A. Sasaki, M. Doebeli, and U. Dieckmann (2013). “Limiting similarity, species packing, and the shape of competition kernels”. In: Journal of theoretical biology 339, pp. 3–13.

Li, Y., R. M. Reeves, X. Wang, Q. Bassat, W. A. Brooks, C. Cohen, et al. (2019). “Global patterns in monthly activity of influenza virus, respiratory syncytial virus, parainfluenza virus, and metapneumovirus: a systematic analysis”. In: The Lancet Global Health 7.8, e1031–e1045.

Lion, S. and S. Gandon (2022). “Evolution of class-structured populations in periodic environments”. In: Evolution 76.8, pp. 1674–1688.

Lion, S., A. Sasaki, and M. Boots (2023). “Extending eco-evolutionary theory with oligomorphic dynamics”. In: Ecology Letters 26, S22–S46.

MacArthur, R. (1970). “Species packing and competitive equilibrium for many species”. In: Theoretical population biology 1.1, pp. 1–11.

Martinez, M. E. (2018). “The calendar of epidemics: Seasonal cycles of infectious diseases”. In: PLoS pathogens 14.11, e1007327.

Martinez-Bakker, M. and B. Helm (2015). “The influence of biological rhythms on host–parasite interactions”. In: Trends in ecology & evolution 30.6, pp. 314–326.

Matsuda, H. and P. A. Abrams (1994). “Runaway evolution to self-extinction under asymmetrical competition”. In: Evolution 48.6, pp. 1764–1772.

Metz, J. A., R. M. Nisbet, and S. A. Geritz (1992). “How should we define ‘fitness’ for general ecological scenarios?” In: Trends in ecology & evolution 7.6, pp. 198–202.

Moriyama, M., W. J. Hugentobler, and A. Iwasaki (2020). “Seasonality of respiratory viral infections”. In: Annual review of virology 7.1, pp. 83–101.

Neumann, G. and Y. Kawaoka (2022). “Seasonality of influenza and other respiratory viruses”. In: EMBO molecular medicine 14.4, e15352.

Nickbakhsh, S., C. Mair, L. Matthews, R. Reeve, P. C. Johnson, F. Thorburn, et al. (2019). “Virus–virus interactions impact the population dynamics of influenza and the common cold”. In: Proceedings of the National Academy of Sciences 116.52, pp. 27142–27150.

Park, J. S. and E. Post (2022). “Seasonal timing on a cyclical Earth: Towards a theoretical framework for the evolution of phenology”. In: PLoS biology 20.12, e3001952.

Pons-Salort, M., M. S. Oberste, M. A. Pallansch, G. R. Abedi, S. Takahashi, B. T. Grenfell, et al. (2018). “The seasonality of nonpolio enteroviruses in the United States: Patterns and drivers”. In: Proceedings of the National Academy of Sciences 115.12, pp. 3078–3083.

Reiner Jr, R. C., M. Geary, P. M. Atkinson, D. L. Smith, and P. W. Gething (2015). “Seasonality of Plasmodium falciparum transmission: a systematic review”. In: Malaria Journal 14.1, p. 343.

Richard, B., A. Qi, and B. D. Fitt (2022). “Control of crop diseases through Integrated Crop Management to deliver climate-smart farming systems for low-and high-input crop production”. In: Plant Pathology 71.1, pp. 187–206.

Rohani, P, C. Green, N. Mantilla-Beniers, and B. T. Grenfell (2003). “Ecological interference between fatal diseases”. In: Nature 422.6934, pp. 885–888.

Sakarchi, J. and R. M. Germain (2025). “MacArthur’s consumer-resource model: a Rosetta Stone for competitive interactions”. In: The American Naturalist 205.3, pp. 306–326.

Sakavara, A., G. Tsirtsis, D. L. Roelke, R. Mancy, and S. Spatharis (2018). “Lumpy species coexistence arises robustly in fluctuating resource environments”. In: Proceedings of the National Academy of Sciences 115.4, pp. 738–743.

Sasaki, A. (1997). “Clumped distribution by neighbourhood competition”. In: Journal of The-oretical Biology 186.4, pp. 415–430.

Sasaki, A. and U. Dieckmann (2011). “Oligomorphic dynamics for analyzing the quantitative genetics of adaptive speciation”. In: Journal of mathematical biology 63.4, pp. 601–635.

Sasaki, A. and S. Ellner (1995). “The evolutionarily stable phenotype distribution in a random environment”. In: Evolution 49.2, pp. 337–350.

Sasaki, A., S. Lion, and M. Boots (2022). “Antigenic escape selects for the evolution of higher pathogen transmission and virulence”. In: Nature ecology & evolution 6.1, pp. 51–62.

Shaman, J., V. E. Pitzer, C. Viboud, B. T. Grenfell, and M. Lipsitch (2010). “Absolute humidity and the seasonal onset of influenza in the continental United States”. In: PLoS biology 8.2, e1000316.

Shirreff, G., S. S. Chaves, L. Coudeville, B. Mengual-Chuliá, A. Mira-Iglesias, J. Puig-Barberá, et al. (2024). “Seasonality and Co-Detection of Respiratory Viral Infections Among Hospitalised Patients Admitted With Acute Respiratory Illness—Valencia Region, Spain, 2010–2021”. In: Influenza and Other Respiratory Viruses 18.10, e70017.

Stroud, J., B. Delory, E. Barnes, J. Chase, L De Meester, J Dieskau, et al. (2024). “Priority effects transcend scales and disciplines in biology”. In: Trends in Ecology & Evolution 39.7, pp. 677–688.

Wainwright, C. E., E. M. Wolkovich, and E. E. Cleland (2012). “Seasonal priority effects: implications for invasion and restoration in a semi-arid system”. In: Journal of Applied Ecology 49.1, pp. 234–241.

Yamamichi, M., A. D. Letten, and S. J. Schreiber (2023). “Eco-evolutionary maintenance of diversity in fluctuating environments”. In: Ecology Letters 26, S152–S167.

