## Supplementary material for "Eco-evolutionary dynamics of pathogen epidemic timing in a seasonal environment"

### Supplementary information for "Eco-evolutionary dynamics of pathogens epidemic timing in a seasonal environment"

#### S1. Derivation of the covariance form of the selection gradient

Here we derive the covariance form of the selection gradient (Eq. (10) in the main text). Since

$$\lambda(\tau(t),\theta) = \beta(\tau(t))\phi(\tau(t),\theta),$$

we have

$$\begin{aligned}\frac{d\lambda(\tau(t),\theta)}{d\theta} &= \beta(\tau(t))\frac{d\phi(\tau(t),\theta)}{d\theta} \\ &= \beta_0(1+\delta E(\tau))\frac{d\phi(\tau(t),\theta)}{d\theta}.\end{aligned}$$

Substituting this into the definition of the selection gradient yields

$$\begin{aligned}\mathcal{S}(\theta) &= \left\langle \frac{d\lambda(\tau,\theta)}{d\theta}\hat{S} \right\rangle \\ &= \beta_0 \left\langle \frac{d\phi(\tau,\theta)}{d\theta}\hat{S} \right\rangle + \delta\beta_0 \left\langle \frac{d\phi(\tau,\theta)}{d\theta}E(\tau)\hat{S} \right\rangle.\end{aligned} \tag{1}$$

Now, define the temporal covariance over one period by

$$\mathrm{Cov}(X,Y) = \langle XY \rangle - \langle X \rangle \langle Y \rangle.$$

Since $\phi$ is normalised over one period ($\int \phi d\tau = 1$), its period integral is independent of $\theta$. Therefore,

$$\left\langle \frac{d\phi}{d\theta} \right\rangle = \int \frac{d\phi}{d\theta} d\tau = \frac{d}{d\theta}\int \phi d\tau = 0.$$

With this, it follows that for any periodic function $X$,

$$\left\langle \frac{d\phi(\tau,\theta)}{d\theta}X \right\rangle = \mathrm{Cov}\left(\frac{d\phi(\tau,\theta)}{d\theta}, X\right).$$

Applying this identity to equation (1), we obtain

$$\mathcal{S}(\theta) = \beta_0 \,\mathrm{Cov}\left(\frac{d\phi(\tau,\theta)}{d\theta}, \hat{S}\right) + \delta\beta_0 \,\mathrm{Cov}\left(\frac{d\phi(\tau,\theta)}{d\theta}, E(\tau)\hat{S}\right),$$

which is the covariance form of the selection gradient.

#### S2. Interpretation of the two covariance terms in the selection gradient

In the main text, we derived the selection gradient for the preferred season $\theta$ and revealed the two key effects underlying the selection: seasonal priority effect and seasonal stabilising effect. To understand the direction of these effects, we analyse the covariance terms that appear in the selection gradient. The function $\phi'(\tau, \theta) = d\phi(\tau, \theta)/d\theta$ represents the sensitivity of transmission success to a shift in $\theta$: it measures how changing $\theta$ locally increases or decreases the infection efficiency around each seasonal time $\tau$. For every $\theta$, the sign of $\phi'(\tau, \theta)$ is determined by whether $\tau$ lies on the clockwise or counter-clockwise side of $\theta$ on the periodic domain (upper panels of Fig. S1). In particular, $\phi'(\tau, \theta)$ is negative when $\tau$ is located on the seasonally earlier side of $\theta$ (the shorter arc from $\tau$ to $\theta$ points forward in time), and positive when $\tau$ is located on the later side (the shorter arc points backward in time).

Each covariance term in the selection gradient takes the form

$$\mathrm{Cov}(W(\tau), \phi'(\tau, \theta)) = \int W(\tau)\, \phi'(\tau, \theta) d\tau$$

where $W(\tau)$ is an epidemiological weighting function and integration is taken over one period. In our case, $W(\tau)$ is either the susceptibility profile $\hat{S}(\tau)$ or the product $E(\tau)\hat{S}(\tau)$. In other words, the sign of the selection gradient is determined by how much weight lies on each side of $\theta$: more weight in the region where $\phi'$ is negative (earlier season) makes the covariance negative, whereas more weight in the region where $\phi'$ is positive (later season) makes it positive.

**Seasonal priority effect** Since the epidemic reduces the host susceptible density, the susceptibility profile $\hat{S}(\tau)$ is relatively large before $\theta$ and relatively small after $\theta$, independent of the value of $\theta$ (middle panels of Fig. S1). Because $\phi'$ is negative in the earlier season and positive in the later season, the overlap between $\hat{S}$ and $\phi'$ always yields

$$\mathrm{Cov}(\hat{S}, \phi') < 0.$$

Biologically, this means that susceptibles accumulate earlier in the season, generating a selection pressure that always pulls $\theta$ toward earlier timing.

**Seasonal stabilising effect** In contrast, the combined factor $E(\tau)\hat{S}(\tau)$ is maximized around $\tau \approx 0.5$ regardless of $\theta$ (bottom panels of Fig. S1). When $\theta < 0.5$, most of the mass of $E\hat{S}$ lies to the later season compared to $\theta$ (where $\phi' > 0$), yielding a positive covariance and pushing $\theta$ later. When $\theta > 0.5$, the dominant mass around $\theta = 0.5$ lies to the the earlier season (where $\phi' < 0$), yielding a negative covariance and pushing $\theta$ earlier. Thus, $\mathrm{Cov}(E\hat{S}, \phi')$ exerts a stabilizing force that pushing $\theta$ toward the seasonal peak of $E\hat{S}$ near $\tau = 0.5$.

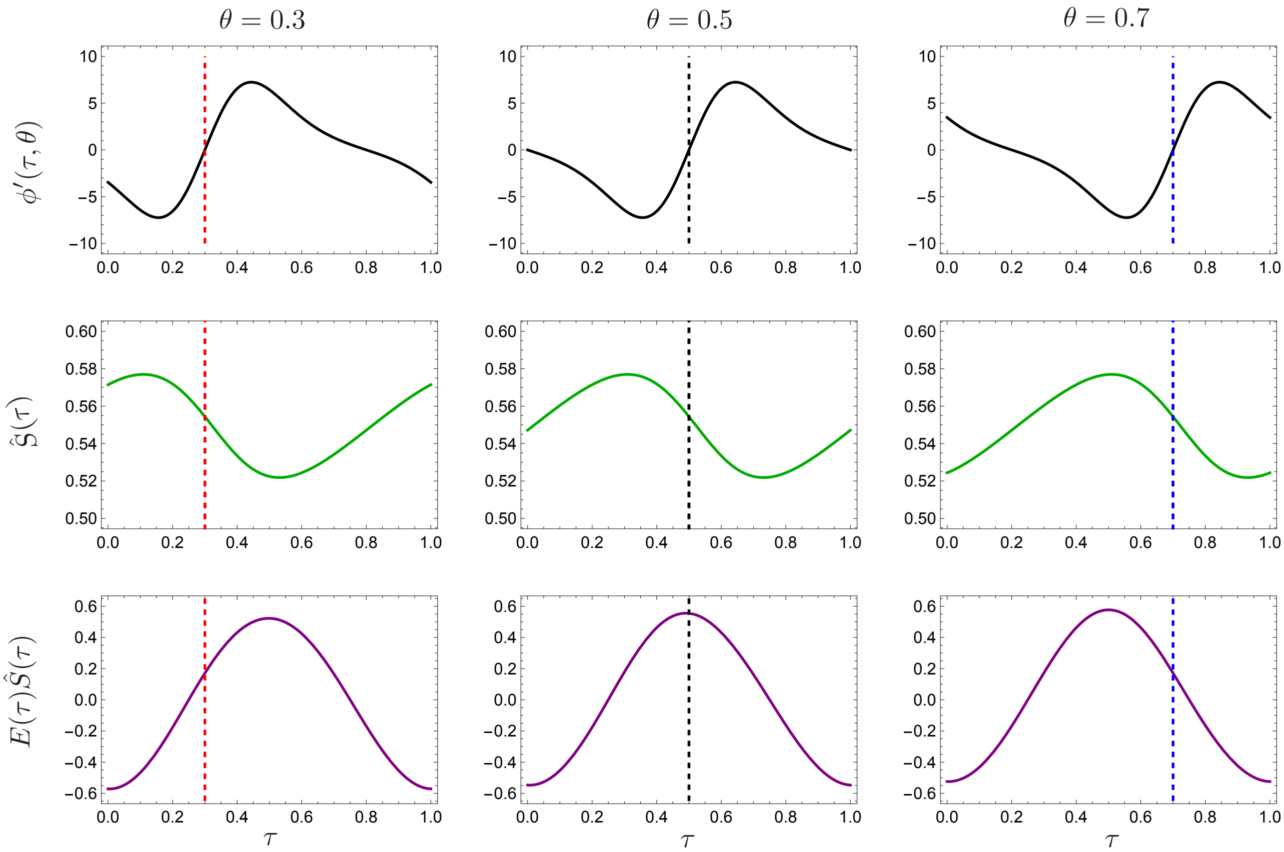


Figure. S1: The components of the covariance terms in the selection gradient of $\theta$. Each row displays, from top to bottom, $\phi(\tau, \theta)$, $\phi'\tau(\tau, \theta)$, and $\phi'\theta(\tau, \theta)$ for three representative values of $\theta$ (0.3, 0.5, 0.7). Colored dashed vertical lines mark the corresponding value of $\theta$. Parameters: $\beta_0 = 2$, $\gamma = 1$, $d = 0.1$, $c = 0.5$, $\kappa = 1$.

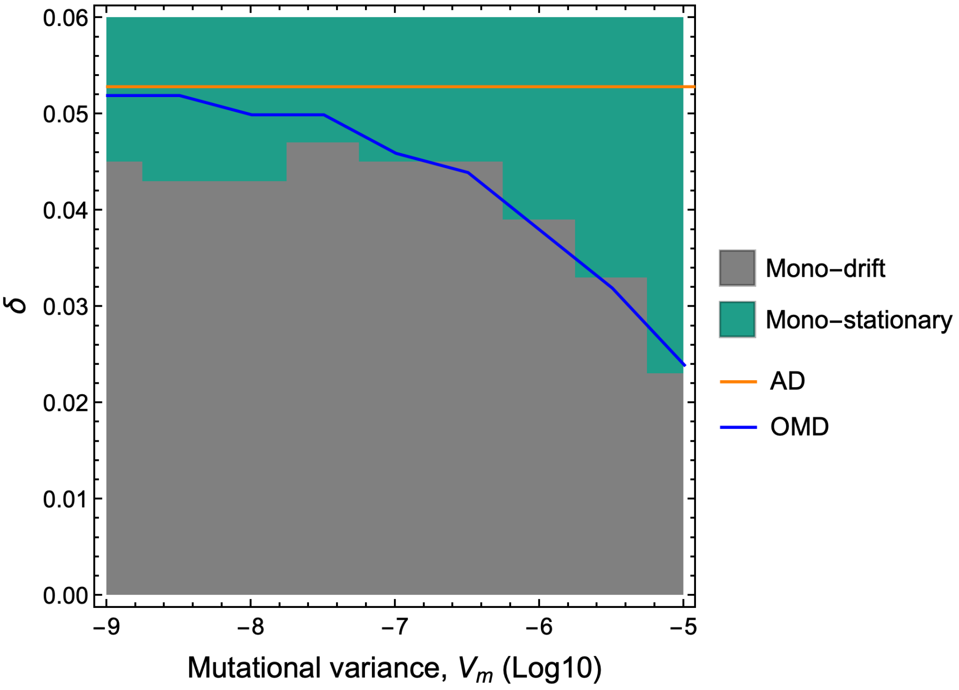


Figure. S2: Variance affects the evolutionary threshold. The boundary between gray and green denotes the critical threshold. Colored area is calculated from evolutionary simulation with the indicated mutant variance. Blue line is obtained from OMD simulations. Parameters: $\beta_0 = 2, \gamma = 1, d = 0.1, c = 0.5, \kappa = 1$

#### S3. Oligomorphic dynamics

Oligomorphic dynamics (OMD) is a recently developed analytical framework that describes the evolution of continuous trait distributions by approximating them with a small number of coexisting morphs (Lion, Sasaki, and Boots, 2023; Sasaki and Dieckmann, 2011). Each morph is represented as a narrow Gaussian distribution in trait space, allowing OMD to extend adaptive dynamics to situations with multiple coexisting phenotypic clusters (Fig. 6A in main text). This approximation yields coupled equations for ecological densities, morph frequencies, and the moments (mean and variance) within each morph. We denote the frequency, the trait mean, and trait variance of morph $i$ by $f_i$, $\bar{\theta}_i$ and $V_i$, respectively. OMD for our system is given by

$$\begin{aligned}
\frac{dS}{dt} &= d - S\sum_i \tilde{\lambda}_i(\tau,\theta_i) f_i I - dS + cR \\
\frac{dI}{dt} &= \left[ S\sum_i \tilde{\lambda}(\tau,\theta_i) f_i - (d+\gamma) \right] I \\
\frac{dR}{dt} &= \gamma I - (d+c)R \\
\frac{df_i}{dt} &= f_i\left(\rho(\bar{\theta}_i) - \bar{\rho}\right) \\
\frac{d\bar{\theta}_i}{dt} &= V_i \frac{\partial \rho(\theta)}{\partial \theta}\bigg|_{\theta=\bar{\theta}_i} \\
\frac{dV_i}{dt} &= V_i^2 \frac{\partial^2 \rho(\theta)}{\partial \theta^2}\bigg|_{\theta=\bar{\theta}_i} + V_M
\end{aligned} \tag{2}$$

where $\tilde{\lambda}(\tau,\theta_i) = \lambda(\tau,\theta_i) + \frac{1}{2}V_i \left.\frac{\partial^2\lambda}{\partial\theta^2}\right|_{\theta=\bar{\theta}_i}$ and $\rho(\theta) = \lambda(\tau,\theta)S - (d+\gamma)$. $V_M$ is the mutational variance (mutation rate times squared mean mutational distance). The first three equations in (2) describe the epidemiological dynamics and the latter three equations in (2) give the evolutionary change of the moments of the trait distribution (i.e. morph frequencies and morph moments). In general, the dynamics of variance depends on the higher moments (skewness and kurtosis, etc) but the assumption of Gaussian distribution simplify the higher moments, providing the closed form of the variance dynamics (moment closure) (Lion, Sasaki, and Boots, 2023; Sasaki and Dieckmann, 2011). OMD requires specifying the (maximum) number of morphs in advance, but by focusing on the moments of the morph, we can determine the actual number of morphs that coexist. By numerically solving (2), OMD predicts the transient evolutionary dynamics of the trait distribution even with polymorphic case, enabling us to characterise the evolutionary patterns.

#### S4. Prediction of evolutionary patterns by OMD

**Numerical simulations:** We numerically solved the OMD system (2) to reproduce the evolutionary patterns observed in the full simulations (Fig. 6B in Main text). We initialized five morphs ($f_i = 1/5$, $V_i = 0.001$) evenly spaced along the trait axis ($\theta_i = i/5$) and ran the OMD for 400,000 years, discarding the first 100,000 as burn-in. The remaining 300,000 years were used to compute two summary metrics: (i) the number of distinct morphs and (ii) the cumulative displacement of the population mean trait $\bar{\theta} = \sum_i f_i\bar{\theta}_i$. Around the boundary between drift and stationary states, evolutionary rates are slow requiring lengthy simulations to determine evolutionary patterns; however, for the majority of parameters, determination of evolutionary patterns is possible within shorter timescales. Note that we start the distribution from $[0, 1]$, but we track the change of $\theta$ on the real axis $(-\infty, \infty)$ to calculate the accumulated moving distance. We run simulations for up to 300,000 years because outcomes are slow to resolve near the boundary between evolutionary drift and stasis, although in much of parameter space the evolutionary regime can be identified in a much shorter time.

**Number of morphs:** Each year, morph $i$ was characterized by its annual mean trait $\langle\theta_i\rangle_t$, frequency $\langle f_i\rangle_t$, and variance $\langle V_i\rangle_t$. For each pair $(i, j)$, we defined a standardized circular distance

$$D_c(i,j) = \frac{\Delta\theta(i,j)}{\sqrt{(\langle V_i\rangle_t + \langle V_j\rangle_t)/2}}$$

where $\Delta\theta(i, j)$ is the shortest circular distance on $[0, 1)$. Morphs with $D_c(i, j) < 2$ were clustered together; otherwise, they were treated as distinct. The number of connected clusters in the resulting adjacency graph was taken as the effective number of morphs and averaged over time. We classified outcomes as one morph ($< 1.005$), two morphs (1.995–2.005), or variable (others). In all cases we analysed, at most three morphs coexisted.

**Directional drift:** To quantify long-term directional change, we computed the annual mean of the population's weighted trait,

$$\langle\bar{\theta}\rangle_t = \left\langle \sum_i f_i\theta_i \right\rangle_t$$

and accumulated interannual differences ,

$$C = \sum_{t=t_a}^{t_z} (\langle\bar{\theta}\rangle_t - \langle\bar{\theta}\rangle_{t-1})$$

As $\theta_i$ evolves on the real axis (not restricted to $[0, 1)$), $C$ represents the total distance travelled in trait space. Drifting systems were defined as those with $C > 3$ (i.e. the mean moves around three times in 30,000 years).

#### S5. Dependency of evolutionary patterns on host life history parameters.

To test the robustness of the OMD predictions shown in Fig. 6, we examined the effects of varying host life-history traits (Fig. S3). All parameter sets exhibited qualitatively similar patterns. A higher host immune loss rate $c$ broadened the range of multiple morphism and increased the tendency towards drift (upper panels in Fig. S3). Similarly, faster host turnover (higher recruitment and mortality rates; larger $d$) made the coexistence of two morphs more likely and slightly increased the tendency towards drift (lower panels in Fig. S3). Notably, the region around the **variable** appears complex; this is because the dynamics in this region exhibit complex patterns, making it numerically unstable and thus sensitive to the effects of numerical criteria.

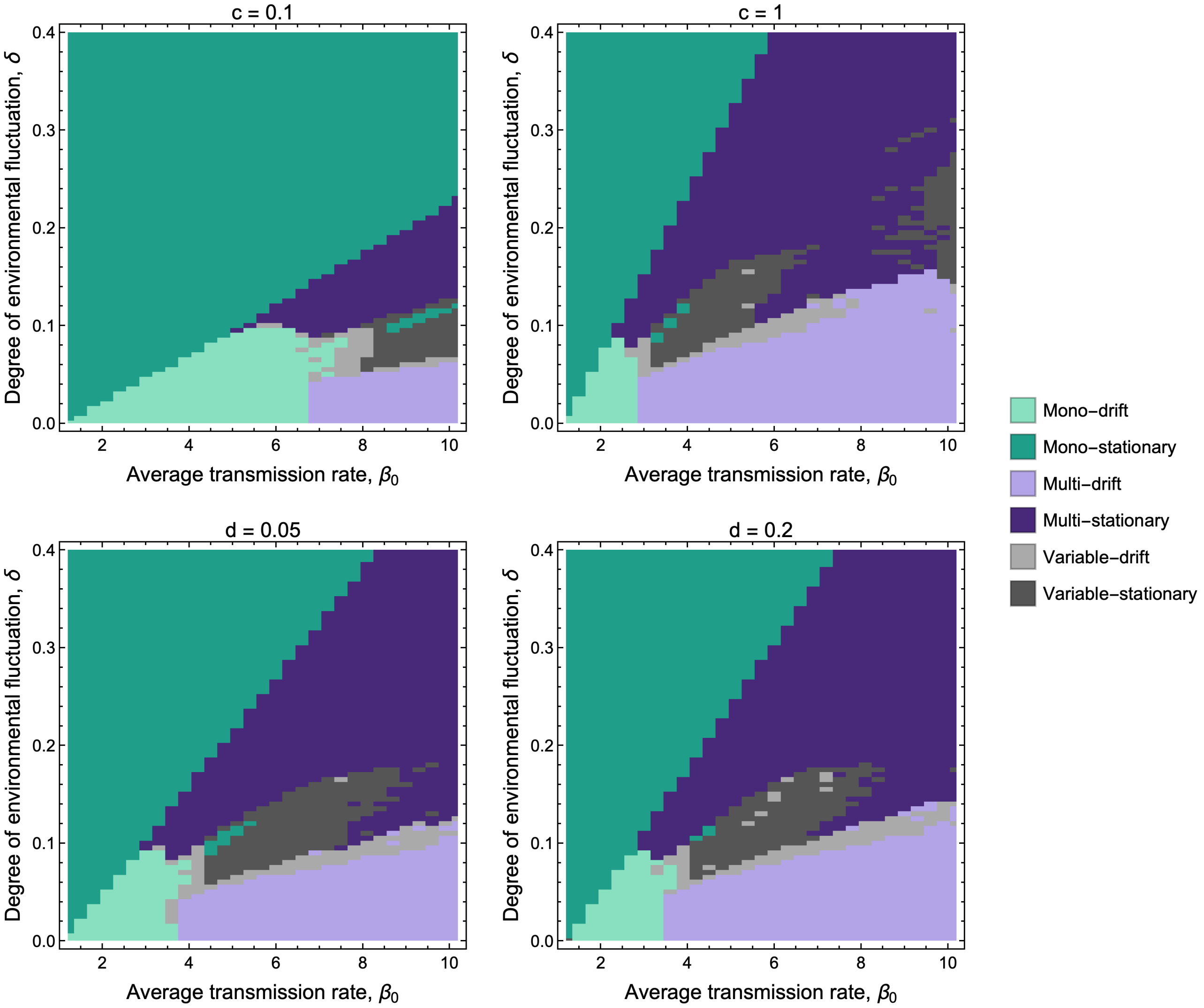


Figure. S3: Evolutionary outcomes predicted by OMD for different parameters. Numerical method is described in Supplementary Information S4. Lighter colours indicate the drifting behaviour, while darker colours indicate that the distribution is fixed or fluctuating around a set of specific seasons. Each panel show the evolutionary patterns predicted by OMD when using the indicated parameter. Default parameter set: $\beta_0 = 2$, $\gamma = 1$, $d = 0.1$, $c = 0.5$, $\kappa = 1$.

### References

Lion, Sébastien, Akira Sasaki, and Mike Boots (2023). “Extending eco-evolutionary theory with oligomorphic dynamics”. In: *Ecology Letters* 26, S22–S46.

Sasaki, Akira and Ulf Dieckmann (2011). “Oligomorphic dynamics for analyzing the quantitative genetics of adaptive speciation”. In: *Journal of mathematical biology* 63.4, pp. 601–635.
